# A low protein diet drives short-and long-term improvements in metabolic health in a mouse model of sleeve gastrectomy

**DOI:** 10.1101/2025.04.23.650310

**Authors:** Julia Illiano, Cara L. Green, Alina Chao, Grace Zhu, Luiz Lopez, Odin Schaepkens, Cholsoon Jang, Dudley W. Lamming, David A. Harris

## Abstract

Despite the largely beneficial impact of bariatric surgery on obesity and metabolic disease, continued post-surgical obesity and weight recurrence is common and may be impacted by diet. While guidelines recommend a high-protein diet based on the theory that this will preserve lean mass, emerging evidence suggests that both humans and mice are metabolically healthier on low protein diets. We assessed the effect of varying dietary protein levels on post-surgical weight loss and weight regain in a mouse model of one type of bariatric surgery, sleeve gastrectomy. We found that a low protein diet optimally drives post-surgical weight loss, boosting energy expenditure and improving blood glucose regulation. Using a multi-omics approach, we identified clusters of differentially expressed genes and metabolites that correlated with these phenotypes and found that diet heavily influences the liver’s molecular response to sleeve gastrectomy. These results suggest that current post-surgical high protein guidelines may limit both the short-and long-term benefits of surgery, and a low protein diet may improve patient outcomes.

## Introduction

Obesity and its resultant metabolic conditions represent an international health crisis^1^ that is expected to impact nearly half of the US adult population by 2030.^2^ Relative to other modern treatments, bariatric surgery remains the most effective and durable therapy to combat obesity and its comorbidities. Owing to the preserved metabolic benefits with reduced long-term risk, sleeve gastrectomy (SG) is now the most popular bariatric surgery worldwide.^3^

Despite its successes, a significant number of patients experience suboptimal weight loss and weight recurrence after surgery.^4,5^ Additionally, while the effects of bariatric surgery on weight loss and metabolism are profound and long-lasting in most, clinically, the maximum weight loss plateaus and most organ improvements occur within the first two years after surgery.^6^ This weight plateau is a highly reproducible phenomenon across multiple randomized and non-randomized studies. Importantly, despite being more metabolically healthy at plateau, patients often remain with obesity.^7–9^ A similar phenomenon is seen with new anti-obesity medications like GLP-1 and GIP single-and dual-agonist therapies, suggesting a shared adaptation between these two approaches to obesity treatment.^10–14^ Thus, it is critical that we identify strategies that further drive the post-surgical weight nadir and bolster the metabolic gains for patients.

The therapeutic potential of post-operative dietary interventions to post-SG physiology is understudied, and discoveries in this space can offer affordable and readily translatable therapies for patients.^15^ Historically, the composition of the post-SG diet has been formulated with the purpose of mitigating potential complications associated with surgery, namely vitamin and mineral deficiencies, hypoalbuminemia, and wound morbidity, which are less prevalent in patients undergoing SG as the absorptive length of the intestine remains unchanged.^16^ Patients are counseled to consume diets high in protein relative to all other macronutrients, with a target of achieving nearly 18-34% of calories from protein (standard recommendation ∼10%).^17^ Further, this is thought to maintain metabolically active lean mass, which decreases after all forms of bariatric surgery.^18^ However, while high protein diets may provide short-term benefits to weight and glucose control given their impact on satiety, they are associated with mid-and long-term diabetes risk, reduced cardiovascular health, cancer, and increased mortality.^19–22^ In fact, more recently, diets that are low in protein but that are not protein insufficient have been shown to induce weight loss and improve metabolic health in mice and humans and extend lifespan in mice.^23–25^ Additionally, the idea of consuming high protein diets to prevent long-term lean muscle mass loss was recently called into question in a study of roughly 3300 twins, which found that high protein diets were associated with sarcopenia while low protein diets were protective.^26^

To date, there have been no comprehensive analyses of varying protein intake on post-SG physiology. Here, we investigated the hypothesis that low protein diets would bolster the metabolic impact of SG in male C57BL/6J mice. Using specially designed, isocaloric diets with low, medium, and high protein concentrations, we rigorously tested the impact of altered post-operative protein intake on weight loss, glucose metabolism, energy expenditure, and the hepatic response to feeding. We found that low protein diets in SG animals produced greater and more sustained weight loss, reduced weight recurrence, improved glucose handling, and induced greater energy expenditure without causing calorie restriction or significant lean mass loss when compared to the other protein groups. We found that in the context of low protein intake, diet is the major driver of the hepatic transcriptomic and metabolomic response to feeding but that with increased levels of protein, there is an increase in the influence of surgery on hepatic physiology. Importantly, we found that SG was protective against the hepatic metabolic damage associated with high protein feeding. A direct comparison of low and high protein post-operative diets in the context of SG revealed that low protein feeding was associated with a large induction of fibroblast growth factor 21 (FGF21), which is a key driver of organismal energy expenditure (EE).^27^

Thus, we conclude that through a low protein post-operative diet, a straightforward and translatable dietary intervention, we can dramatically impact the weight nadir and metabolic health gains following SG in mice. This represents a substantial departure from the status quo and a potential paradigm shift for the management of patients in the weight management clinic.

## Results

### Low protein diet induces maximal weight loss and protects against weight regain following SG

Five-week-old C57BL/6J male mice were fed a high-fat, high sucrose Western Diet (WD; Inotiv TD.88137) for 12 weeks to induce obesity and glucose intolerance. They were then randomized into weight-matched groups to receive either SG or Sham operations. To examine the interaction between post-operative dietary protein consumption and metabolism following SG, both SG and Sham mice were further randomized following surgery to either continue consuming a WD, or to consume a lower-fat diet with either high (TD.22029; 36% calories from protein), medium (TD.180161; 21% calories from protein), or low protein (TD.10192; 7% calories from protein) (**Fig. 1A**; **Table S1**). The three lower-fat diets were isocaloric, with 20% calories from fat and variable carbohydrate to replace any caloric deficit as previously reported.^28^ Body weights and food consumption were tracked longitudinally for 13 weeks following surgery.

**Figure 1.**
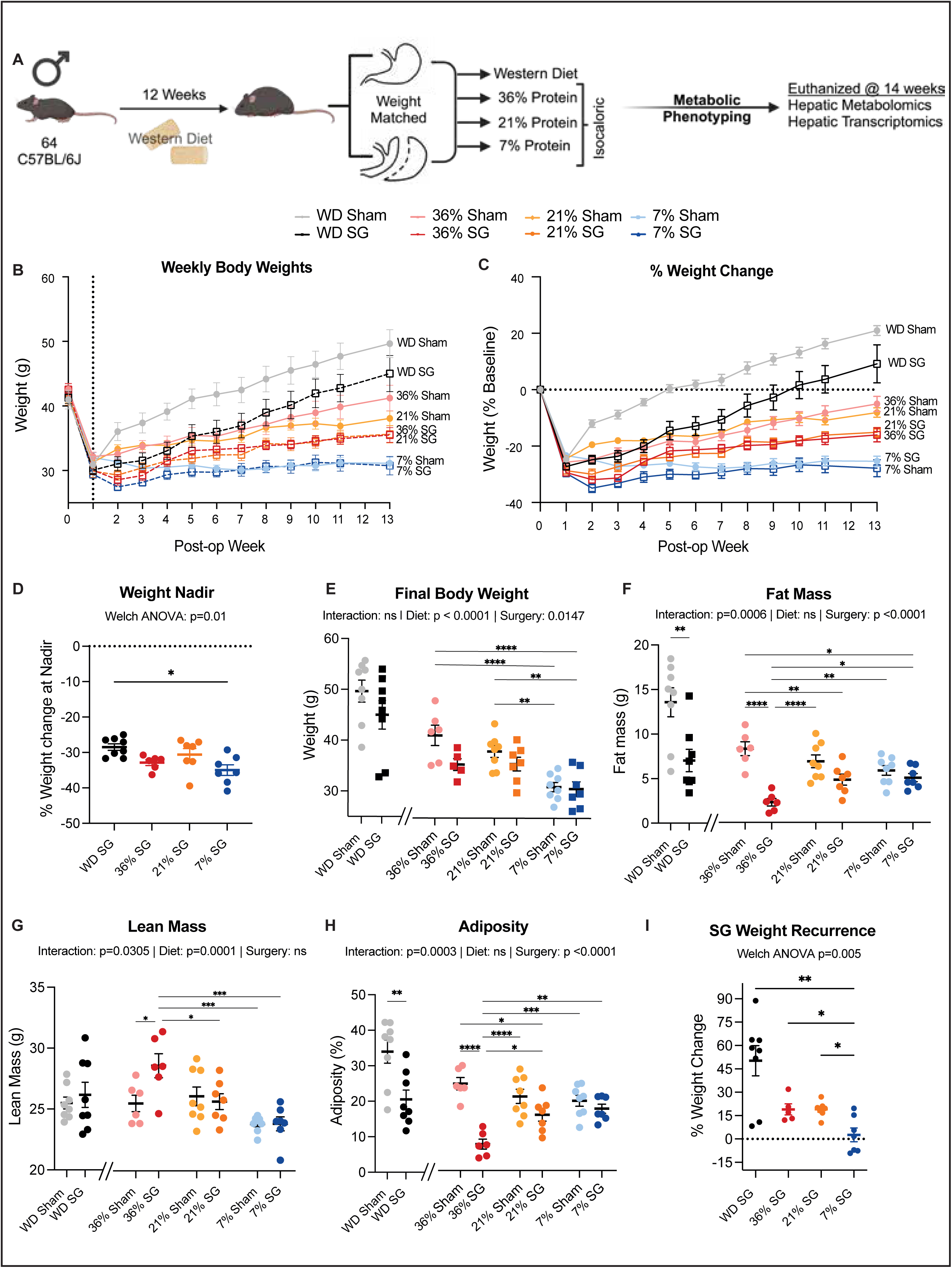
Reduced dietary protein intake following SG maximizes weight loss and maintains lean mass despite increased food consumption. (A) Experimental design. (B-E) total body weight (B), percent body weight change (C), weight nadir (D), and final body weight (E) (F-H) Fat (F) and lean (G) mass was determined by echoMRI and adiposity (H) was calculated. (I) Weight recurrence in SG groups (final body weight as compared to nadir weight) (B-H) n for each measurement is detailed in Table S9. (B, I) SG groups were compared by Welch’s Anova with individual group means compared to 7% SG with Dunnet T3 correction for multiple comparisons. (E-H) WD surgical groups were compared using Welch’s T-test and surgical groups within the protein dietary groups were compared using two-way ANOVA between surgery and diet with post-hoc Tukey corrected test for pairwise comparisons. *p<0.05, **p<0.01; ***p<0.001; ****p<0.0001. (I) Groups were compared to low protein SG using multiple Welch’s T-test with Holm-Šídák method for correction for multiple comparisons.

As we expected, mice in all groups had weight loss during the first postoperative week during Recovery Gel Diet administration. Weight rebounded most rapidly in the WD Sham group, with slower weight regain in WD-fed SG mice (**Fig. 1B**). In mice fed either a high (36%) or medium (21%) protein diet, SG mice weighed less throughout the course of the experiment as compared to Sham mice fed the same diet (**Fig. 1B)**. Intriguingly, SG and Sham mice fed the low (7%) protein diet had identical weight loss and these mice also had the greatest percent weight loss overall (**Figs. 1B-C**). Among mice that underwent SG, those on low protein had the lowest overall weight nadir (**Fig. 1D**). There were effects of both diet and surgery on final body weight, with the lowest final body weight observed in Sham and SG animals fed a low protein diet (**Fig. 1E**).

Body composition analysis was performed during post-operative week 4. In agreement with conventional wisdom regarding the best diet for SG patients to follow post-surgery, we found that SG mice on a high protein diet, despite having the highest overall weight of any SG mice, had the lowest amount of fat mass, the highest amount of lean (fat-free) mass, and the lowest overall adiposity (**Figs. 1F-H**). Notably, this effect was unique to SG mice as high protein-fed Sham mice had significantly more fat and less lean mass than high protein SG mice, which suggests that SG animals uniquely adapt to high protein diets. In fact, when looking across Sham diet groups there was a clear inverse relationship between protein intake and fat mass.

Unexpectedly, there was no relationship between protein intake and grip strength endurance as measured by inverted cling testing (**Fig. S1A**).

We next assessed weight regain (percent change from nadir weight to final weight). As expected, we found that diet heavily influenced weight regain following surgery. Low protein-fed SG mice had the lowest overall recurrence (**Fig. 1I**), with an average of 2.6% weight gain from nadir compared 19.2%, 19.0%, and 50.2 % weight gain found in medium protein, high protein, and WD SG.

### Low protein diets increased carbohydrate fuel utilization and drive energy expenditure in SG mice

We next placed mice in metabolic chambers to examine substrate utilization and organismal energy balance. Both diet and surgery impacted caloric intake, with the highest daily caloric intake generally found in groups with decreasing protein intake (**Fig. 2A**) as previously described.^29^ Thus, changes in weight and body composition in low protein mice was not the result from decreased caloric intake. Interestingly, high protein SG animals had high daily caloric intake compared to high protein Shams, which had the lowest overall food intake of any group.

**Figure 2.**
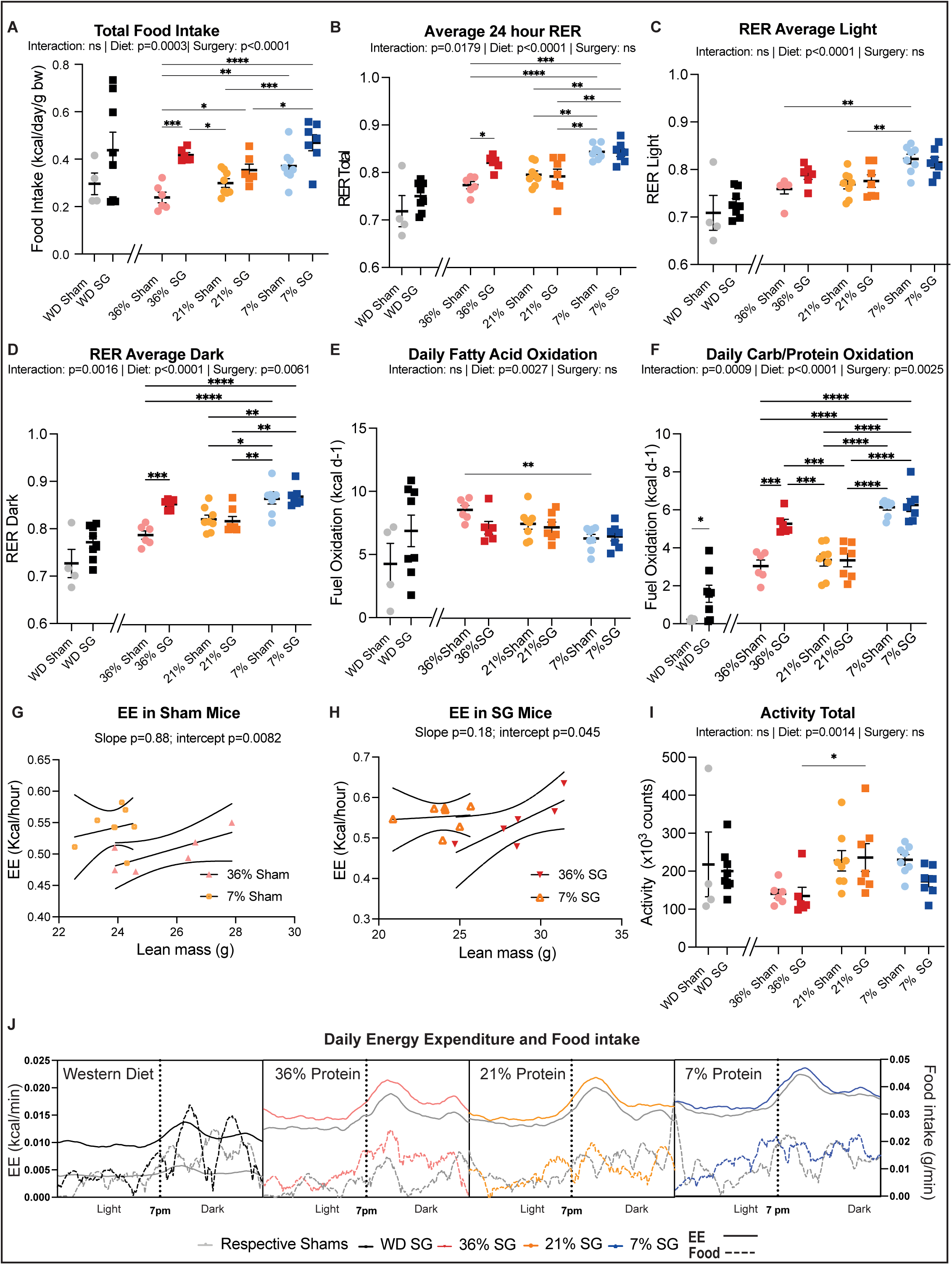
Whole body energy expenditure (EE) was increased in low protein fed animals. A 24-hour metabolic chamber run was used to assess EE, Food intake, Respiratory Exchange Ratio (RER), activity, and fuel oxidation at four weeks post-op. (A) Food intake (**B-D**) The RER across the 24-hour period (B), during the light cycle (fasting; C), and during the dark cycle (feeding; D). (**E-F**) Daily fatty acid oxidation (E) and carbohydrate/protein oxidation were calculated (F) (**G** and **H**) General linear modeling of 24-hour EE normalized to lean mass in high and low protein fed Sham (G) and SG (H) mice. (I) total activity level. (J) Except during low protein feeding, SG animals had higher EE (y-axis; colored trend lines) across a 24-hour period compared to their Sham counterparts. Interestingly, this was most evident during the first half of the dark cycle, where both SG and Sham animals had a spike in initial food intake (x-axis; colored bars) (**A-F; I**) n for each measurement is detailed in Table S9. WD surgical groups were compared using Welch’s T-test and surgical groups within the protein dietary groups were compared using two-way ANOVA between surgery and diet with post-hoc Tukey corrected test for pairwise comparisons. *p<0.05, **p<0.01; ***p<0.001; ****p<0.0001. (**G and H**) General linear modeling with lean mass covariate.

We next examined the respiratory exchange ratio (RER) – the ratio of the oxygen consumed and CO_2_ produced. A value of 0.7 corresponds to lipid use, 0.8 a mixture of fuel sources, and 1.0 corresponds to carbohydrate utilization. RER did not vary within the WD cohort. However, there was a clear impact on RER across the total experimental run (**Fig. 2B**) and during the light phase (fasting; **Fig. 2C**) in the protein groups, with low protein Sham and SG mice experiencing an average RER >0.8. During the feeding phase (**Fig. 2D**), there were clear interactions between surgery and diet on RER. There was a strong trend toward increased and a significantly elevated RER in WD SG and high protein SG mice compared to their respective Shams. Low protein Sham and SG mice had the highest overall RER. Interestingly, there were no differences between high and low protein SG animals. Calculating fatty acid (**Fig. 2E**) and carbohydrate/protein oxidation (**Fig. 2F**)^30,31^ we found no differences in fatty acid oxidation. However, there was both a surgery and dietary effect on daily carbohydrate/protein oxidation, with both low and high protein SG animals with the highest oxidation rate.

Given that low protein-fed SG and Sham animals had reduced weight despite increased food intake, we next investigated organismal energy balance. Correcting for differences in lean mass using analysis of covariance, we found Sham (**Fig. 2G**) and SG (**Fig. 2H**) mice on low protein diet had increased energy expenditure (EE) compared to their counterparts on high protein. There was significantly increased EE in all Sham protein groups compared to WD Shams and a similar trend in SG animals (**Table S2** and **S3**). While ambulatory activity contributes to roughly 10% of an animals’ EE,^32^ there was only a small dietary impact on ambulatory activity, thus not explaining the elevated EE across SG groups. We next mapped EE (solid lines) with food intake (**Fig. 2J**; dotted lines). As expected, EE and food intake increased during the dark cycle for each surgical and dietary intervention group. SG mice in the WD, medium protein, and high protein groups had increased EE compared to Shams, which was not explained by increases in food intake. The differences in EE were most pronounced during the dark (fed) phase. EE did not differ across cycles in low protein-fed mice.

In summary, both SG and low protein diets drove a change in fuel resource utilization and EE, especially in the fed state. However, the impact of low protein diet on EE supersedes the impact of SG alone.

### Low protein diet improves glucose homeostasis independent of insulin sensitivity following SG

To examine the impact of SG and dietary protein on glucose handling, we performed an oral glucose tolerance test (OGTT) two weeks following surgery. Glucose tolerance was dependent upon both diet and surgery status, with SG animals having improved glucose tolerance across all dietary groups, and SG animals on low protein having the lowest AUC (**Figs. 3A-B**). SG animals across all groups also had a lower four-hour fasting blood glucose compared to their respective Sham counterparts, with low protein SG mice having the lowest overall fasting glucose (**Fig. 3C**).

**Figure 3.**
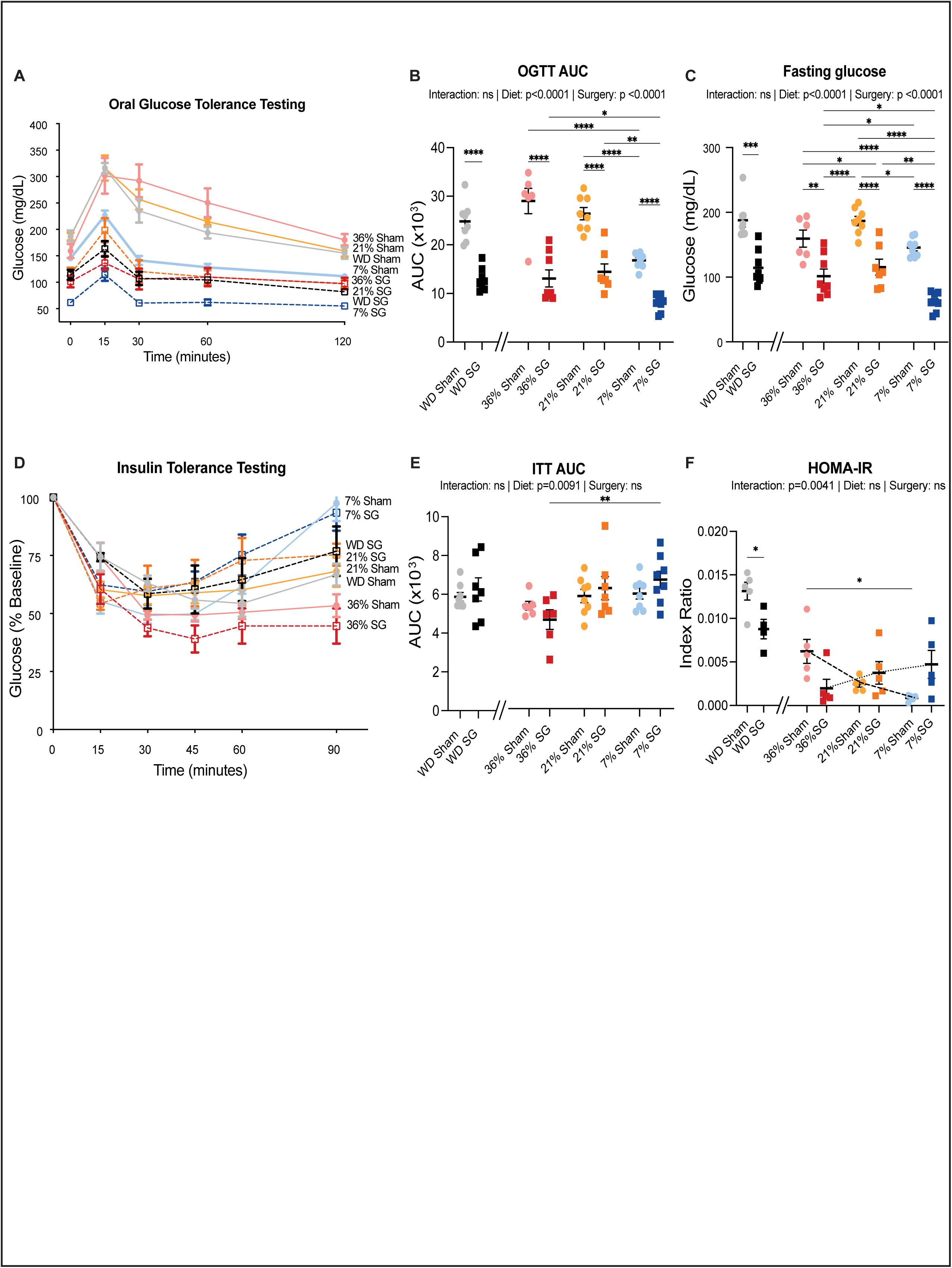
Glucose handling is improved in SG mice on a low protein diet independent of changes in insulin tolerance. (**A and B**) Glucose tolerance testing and calculated area under the curve (AUC). (**C**) Four-hour fasting blood glucose. (**D** and **E**) Insulin tolerance testing and calculated AUC. (**F**) HOMA-IR (**A-F**) n for each measurement is detailed in Table S9. WD surgical groups were compared using Welch’s T-test and surgical groups within the protein dietary groups were compared using two-way ANOVA between surgery and diet with post-hoc Tukey corrected test for pairwise comparisons. *p<0.05, **p<0.01; ***p<0.001; ****p<0.0001.

These changes in glucose tolerance were independent of significant changes in insulin sensitivity as measured by an intraperitoneal insulin tolerance test (ITT) conducted three weeks post-surgery (**Figs. 3D-E**). Surprisingly – and in contrast to previous results by some of us and others – mice on a low protein diet tended to have impaired insulin sensitivity, while mice on a high protein diet tended to have improved insulin sensitivity (**Fig. 3E**). Interestingly, there was no impact on the percent glucose change after 15 minutes (**Fig. S2**), suggesting that there was no impact of surgery or diet on glucose disposal and no worsening of insulin sensitivity in low protein-fed animals. However, in low protein-fed animals there was a rapid return to baseline in the second half of testing, suggesting a strong counter-regulatory process capable of potentiating hepatic gluconeogenesis.^33,34^

As anticipated, WD Sham animals had the highest fasting insulin and Homeostasis Model Assessment of Insulin Resistance (HOMA-IR) across all groups. In Sham animals, there was reduced fasting insulin levels with decreasing levels of dietary protein; however, in SG animals, the opposite was true, with the highest circulating fasting insulin found in WD and low protein-fed SG mice (**Fig. S3**). Thus, SG mice on high protein diet and Shams on low protein diet had the lowest overall HOMA-IR score, suggesting that these two groups had the highest insulin sensitivity.

These results suggest that low protein diets improve glucose sensitivity independent of insulin action but that high protein diets, solely in the context of SG, can impact insulin sensitivity likely through the promotion of lean mass gain.

### Post-SG metabolic phenotype is heavily influenced by protein intake

We used several multivariate analyses to investigate the relationship between protein intake, SG, and metabolic phenotypes. Firstly, we correlated the protein intake of each individual mouse in all three dietary protein groups with 25 phenotypic measurements obtained for each animal in the Sham and SG cohorts. Each correlation was clustered and plotted in a heatmap (**Fig. 4A**). This indicated that, overall, the direction of the relationship between protein intake and metabolic health was similar in both the Sham and SG groups, typically with a stronger association in the Sham group. Protein intake was positively correlated with final body weight, OGTT AUC, and fasting blood glucose, and negatively correlated with calorie intake, EE, RER, and activity levels. Phenotypes in the center of the heatmap were variable between Sham and SG groups, and included fat mass, insulin sensitivity, and inverted cling testing. While protein intake correlated positively with these variables in Sham animals, the relationship was inverted in SG animals. Protein intake appeared to be much more strongly related with lean mass and fasting blood glucose in SG mice, suggesting a potentially more complex relationship with these phenotypes and SG.

**Figure 4.**
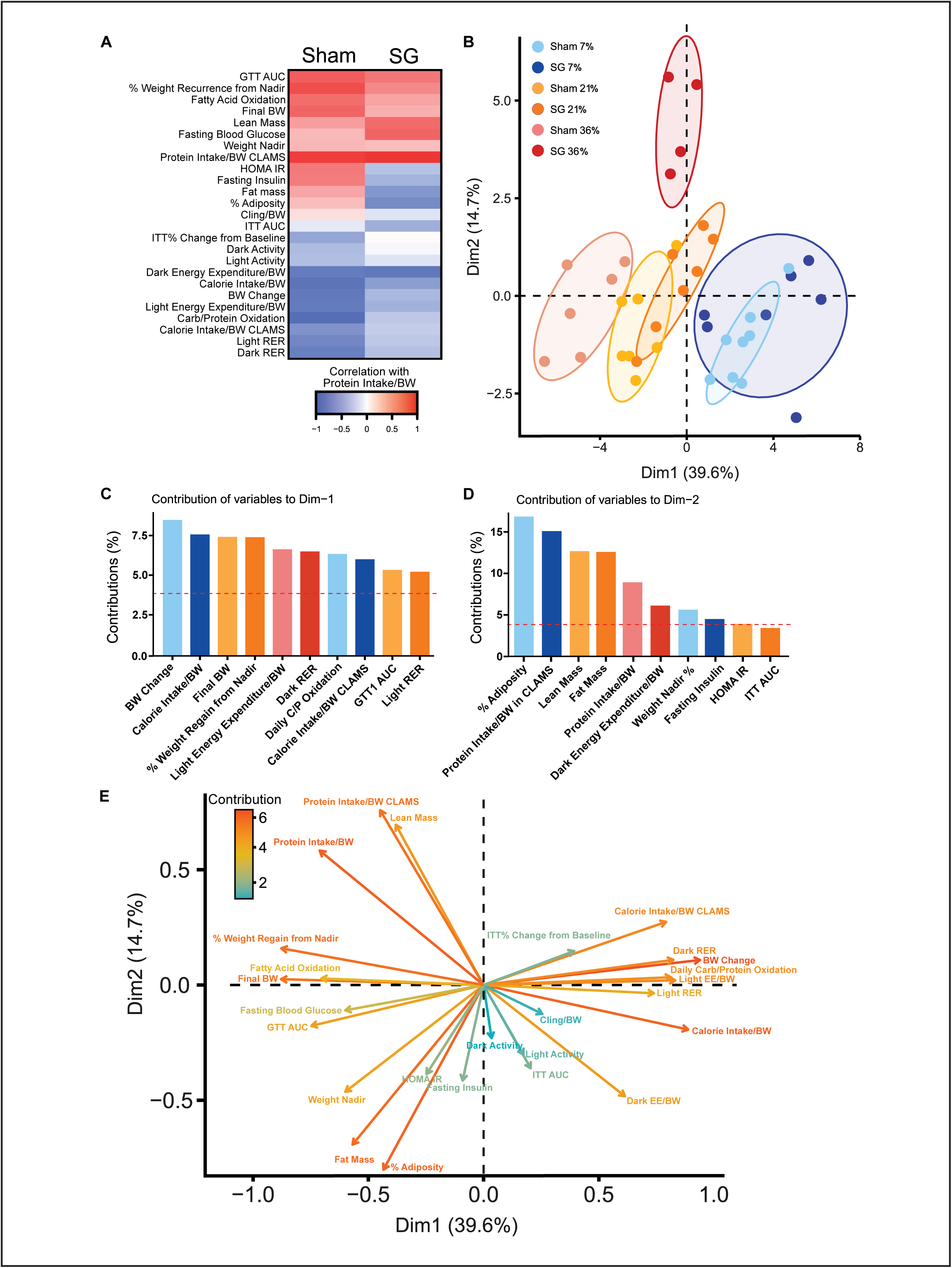
Correlation analysis reveals surgery and diet-specific impacts on physiology. (**A**) Phenotypic measurements were correlated with the protein calorie intake in each mouse (Pearson’s correlations) and clustered via hierarchical clustering. (**B**) PCA unbiased analysis of phenotypic parameters from individual mice. (**C** and **D**) Contribution of each phenotypic measurement to Dimension 1 (C) and Dimension 2 (D) (**E**) Directional visualization of phenotypic parameters. Positively correlated variables have arrows pointing in similar directions and negative correlated variables in opposite directions. Arrow length and color dictate the contribution of each component.

A principal components analysis (PCA) for unsupervised learning was implemented to investigate the relationship between protein intake and surgery across phenotypes. It revealed that with increasing protein intake, there were larger separations between SG and Sham animals (**Fig. 4B**). Dimension 1, which separated groups by protein intake, explained 39.6% of the variance and was most strongly contributed to by body weight change, calorie intake, weight recurrence, and EE (**Fig. 4C**). Dimension 2, which separated out the high protein SG group from the others (**Fig. 4B**), explained 14.7% of variance and was mainly associated with adiposity and protein intake (**Fig. 4D**).

This is highlighted when we explore the contributions of each variable and see that differences between groups were largely driven by changes in protein intake and EE, which contributed most to phenotypic differences by PCA (**Fig. 4E**). Unsurprisingly, we see a positive correlation between protein intake and lean mass but oppositely with EE. These data suggest that diet is a major driver of phenotype when protein intake is low but that SG animals are protected from the detrimental metabolic impact of high protein diets.

### Protein intake heavily influences the hepatic transcriptome and metabolome after SG

Given the critical role the liver plays in regulating metabolism, we conducted a detailed transcriptomic and metabolomic analysis of liver specimens from Sham and SG mice across the three post-operative protein diets in response to feeding. Prior to euthanasia and tissue collection, mice were fasted for 16 hours followed by a 4-hour re-feeding period with their respective diets.

As predicted by the phenotypic similarities between Sham and SG mice on low protein diets, partial least squares-discriminatory analysis revealed that diet had a stronger impact on hepatic transcriptomic variability than SG. When evaluating Dimension 1, a low protein diet clearly induced a different transcriptomic profile than other protein diets. However, SG and Sham mice in the medium and high protein dietary groups were also different (**Fig. 5A**).

**Figure 5.**
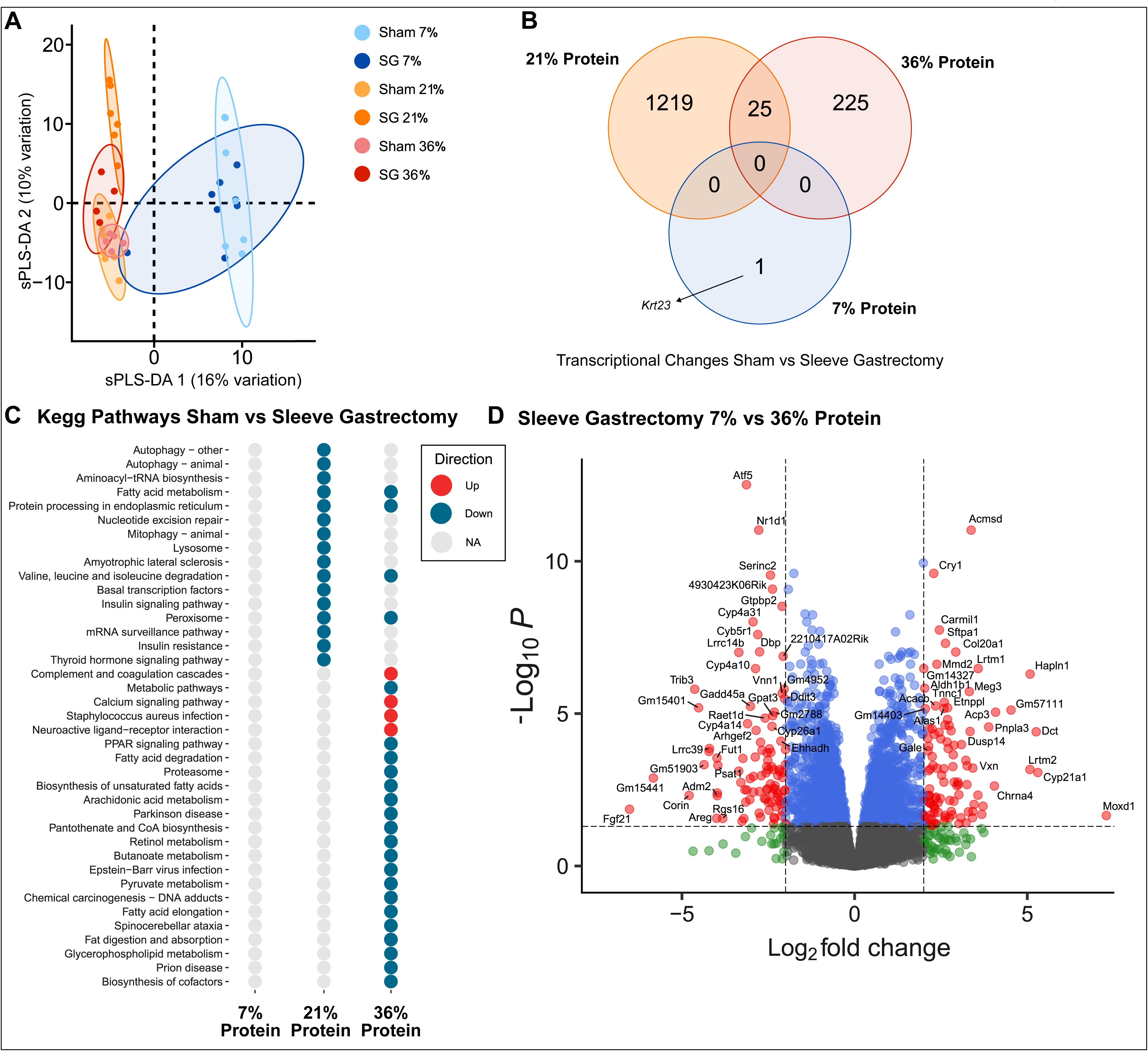
Changes in the post-prandial hepatic transcriptional architecture varied with surgery and diet. (**A** and **B**) Partial least squares-discriminatory analysis of post-prandial whole tissue hepatic transcriptomic data from individual mice (A) and Venn diagram of overlap between differentially expressed genes between conditions (B). (C). Significantly up-and down-regulated pathways for each diet and surgery group were determined via KEGG enrichment. (D). A direct comparison of differentially expressed genes between high and low protein SG mice.

Using an Empirical Bayes linear modelling approach with Benjamini-Hochberg correction, we compared the Sham and SG groups for each diet level. Correspondingly, while there were multiple transcriptomic changes in medium (**Table S4**) and high protein (**Table S5**) SG mice compared to Shams, there were only 25 transcriptomic changes that were shared between SG animals within the high and medium protein diet groups (**Fig. 5B**; **Table S6**). This suggests that each level of protein intake has a unique molecular profile. The largest shared change in medium and high protein SG mice was a 2-fold increase in orosomucoid 3 (*Orm3*), a hepatic acute phase reactant that has an important anti-inflammatory role in hepatic response to stimuli.^35^ There were also shared reductions in *Stat1*, *Irf9*, and *Isg15*, which are key players in interferon signaling.^36–38^ In low protein fed SG and Sham mice, there was only a single differentially expressed gene, keratin 23 (*Krt23*), which was downregulated in SG animals (**Fig. 5B; Table S7**). *Krt23* overexpression has been shown to be associated with hepatic inflammation, failure, and fibrosis.^39^

To identify potential pathways of interest that are enriched in SG animals, we performed a KEGG analysis for each diet group (**Fig. 5C**). Interestingly, there was a clear diet-specific impact of SG on hepatic function, with the most altered pathways occurring within the high protein group. However, animals in the medium and high protein dietary groups had shared reductions in hepatic fatty acid metabolism, endoplasmic reticulum protein processing, branched chain amino acid degradation, and peroxisome activity with SG. There were no substantial pathway-level changes in the low protein group, illustrating the overwhelming impact low protein diet has on hepatic metabolism.

A direct comparison of SG animals in the low and high protein groups reveals a massive change in the hepatic transcriptome (**Fig. 5D**). Most notably, there was a 6.5-log2 fold reduction in the expression of fibroblast growth factor 21 (*Fgf21*) in high protein SG animals (**Table S8**). Hepatic FGF21 is a known driver of EE, insulin sensitivity, and weight loss in mice and humans on either low protein or low isoleucine diets.^40,41^ Additionally, there were multiple upregulated gene targets in high protein SG animals – monooxygenase DBH1 (*Moxd1*)^42,43^, hyaluronan and proteoglycan link protein 1 (*Hapln1*), collagen type XX alpha 1 chain (*Col20a1*), cholinergic receptor nicotinic alpha 4 subunit (*Chrna4*)^44^, patatin-like phospholipase domain-containing 3 (*Pnpla3*),^45^ and aminocarboxymuconate semialdehyde decarboxylase (*Acmsd*)^46^ – which are associated with hepatic fibrosis, extracellular matrix remodeling, and metabolic dysfunction-associated liver disease (MASLD; formerly non-alcoholic fatty liver disease) progression. *Acmsd* plays a pivotal role in the hepatic *de novo* synthesis of NAD+ from tryptophan, whereby it shuttles precursors toward acetyl-coa production and entry into the TCA cycle as opposed to toward NAD+ production. This imbalance increases DNA damage and worsens MASLD,^46^ thus suggesting that SG animals on high protein diet have increased hepatic remodeling compared to those on low protein.

We also performed untargeted metabolomics to investigate significant changes in livers across dietary and surgical groups. As expected, the largest separation between SG and Sham animals was seen in the high protein dietary group, while there was near-complete overlap in the low protein group (**Fig. 6A**). While SG induced significant changes in individual metabolites within each dietary group, there were no individual metabolites that changed significantly across all three protein groups in response to SG and only two metabolites varied in a similar direction.

**Figure 6.**
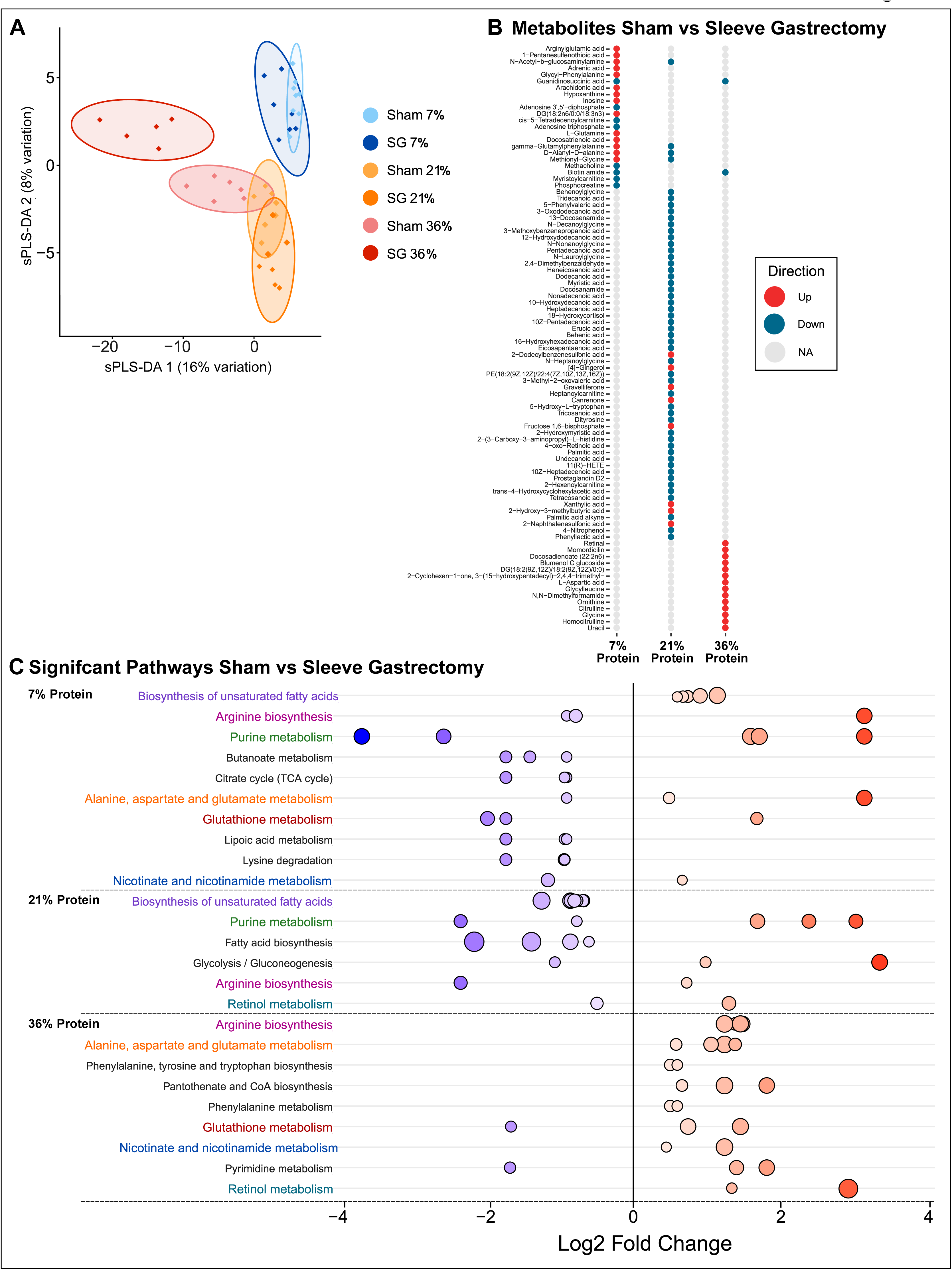
Changes in post-prandial hepatic metabolic pathways varied with surgery and diet. (**A**) PCA of post-prandial whole tissue hepatic metabolomic data from individual mice. (**B**) Differences in individual metabolites from SG and Sham animals across dietary groups. (**C**) Significantly up-and down-regulated metabolic pathways were determined using KEGG enrichment analysis. Dots represent individual pathways, and color indicates log_2_ fold change. Matching pathways between groups are bolded and highlighted.

Thus, the largest driver of the hepatic metabolite composition (**Fig. 6B**) in the post-prandial state was diet. SG in both the low and high protein context led to a decrease in guanidinosuccinic acid, which is a product of Arginine transamination and elevated in uremia associated with high protein feeding.^47–49^ There was an additional decrease in phosphocreatine in low protein SG animals suggesting a complete reduction in this pathway.

Interestingly, the only shared change across all three protein groups was in “arginine biosynthesis” with both low and high protein SG animals having an increase compared to their respective Shams (**Fig. 6C**). Low protein SG animals also had perturbations in purine metabolism driven by an increase in inosine and hypoxanthine and a reduction in both adenosine 3’,5’-diphosphate and adenosine triphosphate, suggesting a favoring of the deamination of adenosine in this context **(Fig. 6B**, **6C**). Inosine has been shown to suppress inflammatory responses across multiple organs and is protective against endothelial dysfuntion.^50^ Lastly, in the high protein context, SG induced increases in amino acid, retinol, and glutathione metabolism.

Hepatocytes and hepatic stellate cells are the dominant locations for retinol storage. Others have shown that decreasing hepatic retinol stores are associated with worsening liver disease and retinol catabolism regulates hepatic lipid stores.^51^ The elevation in glutathione metabolism suggest that high protein SG animals are more protected against redox imbalance associated with high protein feeding.^52^

### Data integration

To define the relationship between hepatic molecular changes, organismal phenotypic changes, and metabolism, we employed two integration methods. Initially we used weighted gene co-expression network analysis (WGCNA; **Fig. 7A**).^53^ We quantified associations of individual genes with traits of interest: protein intake (brown) and EE (turquoise). In mice that had undergone SG, there was a strong correlation between protein intake and the brown module of genes (**Fig. 7B**). These genes were enriched for protein targeting to the endoplasmic reticulum and Golgi vesicle transport, suggesting that changing protein intake may affect intrinsic amino acid processing. Several phenotypes correlated strongly to the turquoise module of genes, including positively with EE and negatively with fasting blood glucose, which is associated with fatty acid oxidation (**Fig. 7C**). WGCNA analysis of the Sham mice rendered fewer unique modules of interest (**Fig. S4**), suggesting a unique relationship between protein intake and surgery at the module-trait level.

**Figure 7.**
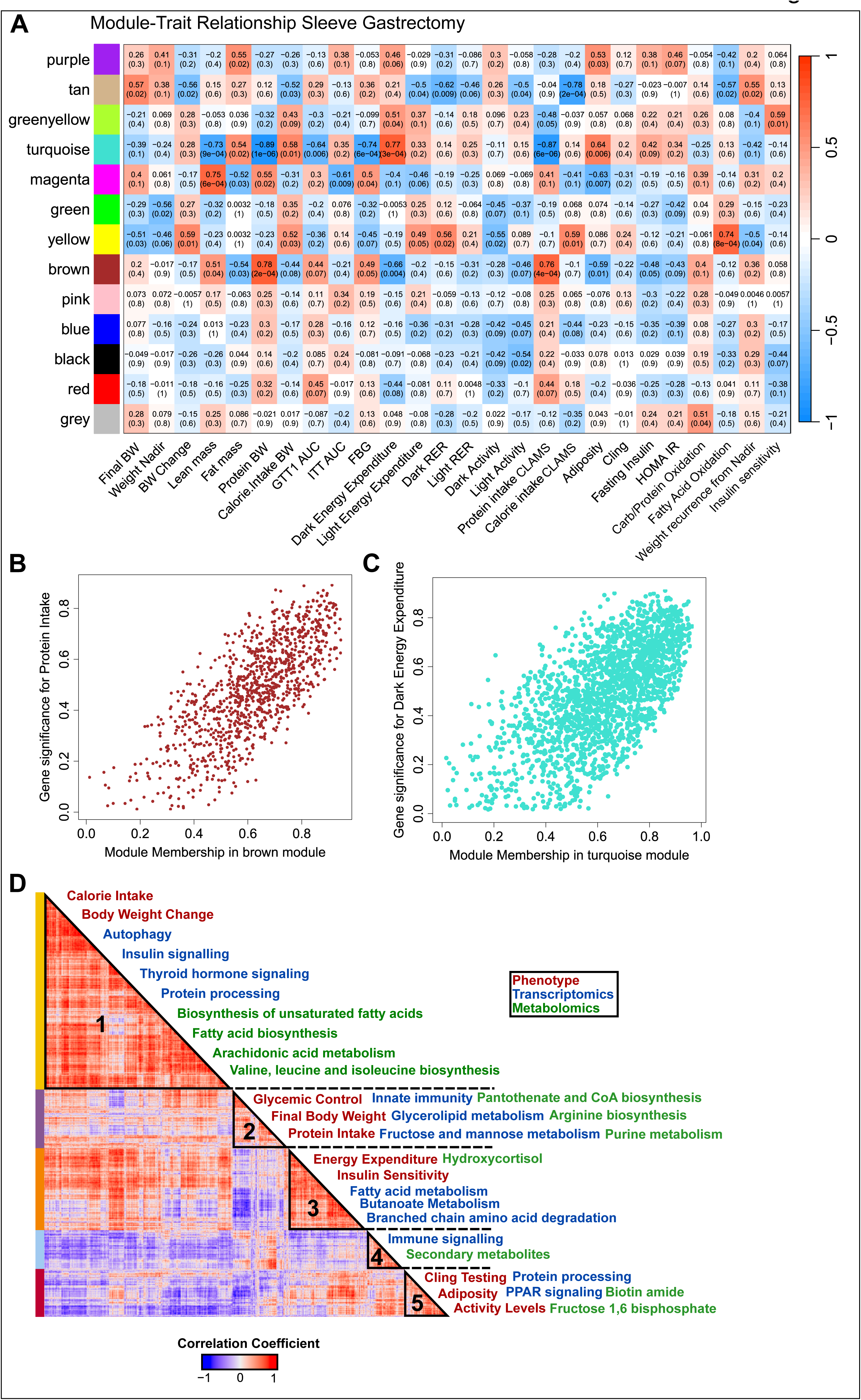
Body weight and EE correlate with branched chain amino acid catabolism. (**A**) Weighted gene co-expression network analysis of differentially expressed genes across SG groups. (**B** and **C**) Protein intake and EE correlated most strongly with the brown (B) and turquoise (C) modules, respectively. (**D**) Integration of phenotypic, transcriptomic, and metabolomic data across individual mice resulted in five distinct modules.

In addition to WGCNA, we integrated phenotypic, transcriptomic, and metabolomic data as done previously.^28^ In brief, we first identified the statistically significant hepatic transcriptomic and metabolomic changes induced by SG across each dietary protein group. Next, we concatenated these changes with the phenotypic data from each individual mouse, which results in a total of 1607 inputs. Pairwise comparisons were made and were rendered by hierarchical clustering (**Fig. 7D**). This resulted in five modules of significant changes containing phenotypes, metabolites (which we enriched for pathways using Metabonanalyst), and genes (which we enriched using the KEGG database to give pathway context to individual gene changes).

As expected, we found that valine, leucine, isoleucine (branched-chain amino acids), and fatty acid biosynthesis was correlated with insulin signaling, protein processing, and body weight change (**Fig. 7D**; cluster1). Additionally, branched chain amino acid degradation and fatty acid metabolism was correlated with EE (**Fig. 7D**; cluster3), as expected by previous work.^40,54^ The correlation of EE with butanoate metabolism signifies a potential role of microbial-derived short chain fatty acids to whole body metabolism. Butyrate predominately comes from the intestinal microbial fermentation and serves as a primary fuel source for the intestine. Importantly, it also has a wide-ranging impact on metabolism through actions upon hepatic, central nervous, and brown adipose tissues.^55–57^ Butyrate supplementation has been shown to limit accretion of fat mass, improve glucose homeostasis, and increase EE in mice on high fat diet.^56^ Additionally, gut-derived butyrate plays a key role in the gut-brain axis and in the regulation of neuronal metabolism.^55,57^

Several phenotypes associated with glycemic control correlated with gene expression related to innate immunity, glycerolipid metabolism, and fructose mannose metabolism, as well as the biosynthesis of Arginine and purine metabolism (**Fig. 7D**; cluster 2).

## Discussion

Bariatric surgery is a highly effective intervention for obesity and metabolic disease; however, some patients do not lose as much weight as desired, and weight recurrence is common.^7,58–61^ Patients experience a reliable weight nadir and plateau roughly two years from the index operation.^62^ These factors remain a significant barrier to the successful treatment of obesity and the fear of suboptimal response and recurrence remain major drivers of poor patient referral for treatment.

We set out to define how post-operative diet influences the initial and long-term success of bariatric surgery in a mouse model of SG. Dietary macronutrient composition is a major contributor to health and disease. However, the current consensus guidelines in post-bariatric surgical nutrition lack rigorous analysis. Dietary protein has been shown to regulate host metabolism. Current bariatric surgical guidelines recommend that patients consume an increase in dietary protein ranging from 1.1 to 2.1 g/Kg of ideal body weight (18% to 34% of calories from protein) following SG and other forms of bariatric surgery.^17,63^ For periods of “active weight loss,” it is recommended to consume protein intake along the higher range in order to maintain lean mass and promote satiety.^63^ However, in a study of 24 humans that underwent SG, there was no correlation (R^2^=0.154; p = 0.064) between protein intake and lean mass loss after surgery, and the importance of lean mass preservation to metabolic health following SG has not been substantiated.^18^

Here, we have performed the most comprehensive analysis of the impact of post-operative protein intake on the metabolic health of mice following SG. Broadly, we have shown that altering post-operative protein nutrition dramatically impacts host metabolism in the setting of SG. The phenotypic and hepatic molecular response to SG was highly diet dependent, with SG having the greatest influence on metabolism in the context of high and medium protein diets. SG protected these mice from weight gain, fat accretion, and abnormal glucose homeostasis, which is known to be associated with high protein diet feeding.^22,28,40,64,65^ In fact, for Sham animals, high protein feeding was equally or more detrimental as WD feeding to oral glucose tolerance (**Fig. 2B**) and fasting glucose levels (**Fig. 2C**). In these dietary arms, SG led to a reduction in adiposity and higher EE. In the high protein arm, SG also led to a clear robust increase in lean mass with an associated trend toward improved insulin sensitivity. The SG hepatic transcriptome and metabolome revealed that SG appeared protective against pro-inflammatory and oxidative states that are typical of high protein feeding.

The dominant contribution to metabolic health in the setting of low protein intake was diet, which was unexpected. While there was some added benefit to systemic glucose control in SG animals on low protein diet, restricting protein intake to 7% total calories improved weight loss, increased EE, and drove hepatic metabolism irrespective of surgical randomization. This was particularly evident when assessing overall change in the hepatic transcriptome (**Fig. 5A**) and metabolome (**Fig. 6A**) in response to feeding. The lower the protein intake, the higher the overlap between Sham and SG mice. While there were some metabolite changes specific to SG animals on a low protein diet, there was only a single gene-level change when compared to low protein Shams. How this would differ in the fasted state or across other organs remains unclear.

Perhaps the most clinically relevant comparison, however, is between SG mice on high and low post-operative protein. In a direct contradiction to the current dietary dogma for patients in the weight loss clinic, low protein-fed SG mice had 2-fold more weight loss, higher EE per gram of lean mass, a 6.5-fold higher expression of FGF21, 16.4% reduction in weight recurrence, and a 40% reduction in fasting glucose, representing a highly significant metabolic benefit to low protein diets.

The induction of FGF21 may be a major contributor to these benefits as it has been well established that benefits of low protein and specific amino acid diets are dependent upon the induction of hepatic FGF21. FGF21 is an insulin-sensitizing hormone produced by the liver in response to nutrient stress and is highly induced by low protein diets.^66,67^ This leads to increases in EE, reduced body weight gain, heightened glucose sensitivity, altered hepatic metabolism, and lifespan extension, at least in C57BL/6J male mice.^28,68,69^ The impact of bariatric surgery on FGF21 levels in humans is unclear with some studies reporting an increase, no change, or a decrease in circulating levels.^70,71^ However, in a study of Roux-en-Y gastric bypass (RYGB), an alternative form of bariatric surgery, whole body FGF21 knock out did not impact weight loss, fat mass, glycemic control, or EE,^71^ thus further highlighting that the induction of FGF21 in our low protein animals is likely diet driven and a major contributor to the benefits seen in the low protein SG mice.

Interestingly, we found that body weight and EE clustered with insulin sensitivity and branched chain amino acid (BCAAs; valine, leucine, and isoleucine) catabolism (**Fig. 7D**). Both SG and RYGB have been shown to reduce circulating BCAAs.^72–76^ In the context of RYGB, this reduction is mediated by the action of hepatic FGF21 to reduced hepatic BCAA breakdown.^77^ However, this action does not appear to be critical as increasing BCAA supplementation and ablation of BCAA breakdown through the genetic ablation of Pp2Cm did not curb the metabolic improvements of SG in mice.^78^ Interestingly, we found that high and low protein feeding was associated with increased hepatic glutathione production in SG mice, which has been previously shown to be directly linked to the shuttling of BCAA catabolic nitrogen-containing products from brown adipocytes to the liver, leading to reduced oxidative stress and improve hepatic insulin sensitivity.^79^

In addition to the role BCAAs play in FGF21 production, BCAAs are also known to play an outsized role in the host maintenance of metabolically active lean mass, and bariatric surgery and other metabolic therapies like incretin mimetics are known to reduce both fat and fat-free mass.^80^ In our study, high protein feeding increased fat-free mass, which is in keeping with a study of humans following SG where post-operative BCAA and Vitamin D supplementation increased fat-free mass compared to whey protein supplementation alone. This occurred without dramatically impacting total weight loss. However, unlike the standard whey protein group, the BCAA and Vitamin D supplemented group did not have improvement in their hemoglobin A1c across the study.^80^ Importantly, we found that low protein feeding was not associated with lean mass loss when compared to animals on medium protein diet, and low protein SG animals were more metabolically healthy than high protein SG animals.

In aggregate, our results show that a dietary intervention can have a major impact on host global and hepatic physiology following SG. Contrary to standard practice, we found that low protein diets could further improve metabolic health of SG mice without causing significant lean mass or strength loss. These findings may have major clinical implications and suggest that the dogmatic march toward higher and higher protein diets as a strategy for weight loss and weight maintenance following SG is unfounded. In fact, the only groups that enjoyed freedom from any degree of weight recurrence, the Achilles’ heel of bariatric surgery, were those on a low protein diet, perhaps informing the long-term implications of this post-operative dietary strategy.

## Limitations

Limitation of our work include the relatively short length of study, the use of young mice, and the reliance on a single sex and strain. Green and colleagues showed that sex and strain influence the response of mice to low protein diets and the relative importance of FGF21 to these effects.^28^ Future studies are needed to define how sex, strain, and age influence our findings.

Further research is also needed to define the exact level of protein intake and optimal amino acid composition necessary to maximize the impact on EE while minimizing any potential harms. It is also unclear if the dietary intervention could be minimized to the first post-operative year during the steepest of the metabolic health gain or whether patients will need to be on a long-term low protein dietary program.

One important factor to consider is how exercise will potentiate the impact of diet. While resistance training did not cause a differential impact on strength in a study of high and low protein-fed mice, it was protective against fat accretion in the context of high protein diets and improved insulin sensitivity.^81^ Future studies will need to evaluate how resistance training impacts our findings, especially given the increase in lean mass seen in our high protein SG mice.

Lastly, it is important to remember that different bariatric surgeries, while linked together in most clinical studies, are not a monolith and lead to different physiologic outcomes. As such, the impact of low protein diets in the context of RYGB will need to be independently evaluated.

## Acknowledgments

We thank all members of the Harris (WiSLiM), Lamming, and Jang lab for their feedback and Drs. Galmozzi and Alexander for their insightful discussions. Dr. Harris is supported by the UW Department of Surgery, School of Medicine and Public Health, Wisconsin Alumni Research Fund, and the Office of the Vice Chancellor for Research. Additionally, Dr, Harris has funding through the Washington University Diabetes Research Center (P30DK020579), Wisconsin Alzheimer’s Disease Research Center (P30-AG062715), and a grant from the Wisconsin Partnership Program at the UW School of Medicine and Public Health (ID 6770-2024). We would like to thank Medtronic (Minnepolis, MN) for a grant allowing the purchase of linear cutting staplers.

The Lamming lab is supported in part by the NIA (AG056771, AG081482, AG084156, and AG085898), the NIDDK (DK125859), the Wisconsin Partnership Program, and startup funds from UW-Madison. Dr. Green was supported in part by Dalio Philanthropies, a Glenn Foundation Postdoctoral Fellowship, and by a Hevolution Foundation award (HF-AGE AGE-009). The Lamming lab was supported in part by the U.S. Department of Veterans Affairs (I01-BX004031), and this work was supported using facilities and resources from the William S. Middleton Memorial Veterans Hospital.

The Jang lab is supported by R01-AA029124 (NIAAA), R21-AA030358, and the Pew Foundation

The content is solely the responsibility of the authors and does not necessarily represent the official views of the NIH. This work does not represent the views of, the Department of Veterans Affairs, or the United States Government.

## Author Contributions

JI, DLW, and DAH conceived of and designed the experiments. JI, AC, GZ, LL, OS, CJ, and DAH performed the experiments. AC and CJ performed metabolomics. CG performed all bioinformatics work. JI, CH, AC, CJ, DWL, and DAH analyzed the data. JI, DWL, and DAH wrote the manuscript.

## Materials and Methods

### Animals

Male C57BL/6J mice were purchased from Jackson Laboratories at five weeks of age and preconditioned on a high-fat WD (Inotiv TD.88137; 42% calories from fat; 15% calories from protein; 43% calories from carbohydrates) for 12 weeks to induce obesity and glucose intolerance. Mice were housed in groups of two and had ad libitum access to food and water in a climate-controlled environment with a 12-hour light/dark cycle. All procedures were approved by the Institutional Animal Care and Use Committee (UW-Madison and William S. Middleton Memorial Veteran’s Hospital), and animals were cared for according to guidelines set forth by the American Association for Laboratory Animal Science in an AALAC-accredited facility.

### Sleeve gastrectomy and sham procedures

At 17 weeks of age, mice were weight matched and randomized into either a Sham or SG surgical group as previously described.^82^ In brief, mice were placed on a recovery gel diet from 48 hours prior to surgery to 6 days post-surgery before going on their respective diets (Clear H_2_0, Westbrook, ME). Mice were anesthetized with isoflurane under sterile conditions. At the time of surgery, mice received weight-based saline, norocillin, and buprenorphine (Ethiqua XR; Fidelis Animal Health, North Brunswick, NJ). SG consisted of a midline laparotomy, short vessel ligation, and removal of the entire non-glandular stomach using a linear cutting stapler (Medtronic, Minneapolis, MN). Sham surgery similarly consisted of a laparotomy, short vessel ligation, and manipulation of the stomach along the would-be staple line.

### Diets

On the morning of post-operative day 7, mice were equally placed into one of four diet groups: continued WD (TD.88137; 42% calories from fat, 42.7% calories from carbohydrates, 15.2% calories from protein) or one of three isocaloric, natural-sourced protein diets (**Table S1**). The three protein diet groups were as follows: high protein (Inotiv, TD.22029; 36% calories from protein), medium protein (Inotiv, TD.180161; 21% calories from protein), and low protein (Inotiv, TD.10192; 7% calories from protein). The carbohydrate percentage was altered to account for the differences in calories from protein.

### Functional glucose testing

Body weights and food consumption were tracked longitudinally. OGTT and ITT were performed after a four-hour fast (7 am to 11 am). The OGTT was performed on postoperative week 2. Mice received an oral bolus of 30% glucose (1g/kg) and blood glucose was measured at time 0, 15, 30, 60, and 120 minutes. The ITT was performed on postoperative week 3 and mice received an intraperitoneal bolus of regular insulin (0.5 u/kg; Humulin R: Eli Lilly USA). Blood glucose was measured 0, 15, 30, 60, and 90 minutes. The blood glucose measurements were collected using a Bayer Contour glucose meter and test strips. Glucose Stimulated Insulin Secretion (GSIS) was performed after a four-hour fast and 10 ul of plasma was collected at fasting and 15 minutes after receiving an oral gavage of 30% glucose (2g/kg). Circulating insulin was measured using Crystal Chem’s Ultra-Sensitive Mouse Insulin ELISA Kit (Elk Grove Village, IL).

### Body composition and indirect calorimetry analysis

Body composition was assessed using the EchoMRI Body Composition Analyzer. Indirect calorimetry was performed using Oxymax/CLAMs metabolic chamber system. Mice were individually housed in CLAMs cages at 22°C and measurements for CO_2_ production, O_2_ consumption, heat production, RER, ambulatory activity, and food intake were recorded for each mouse. On postoperative week 4, mice were placed in CLAMs cages for ∼48 hours with a 12-hour light and dark cycle. The first 24 hours were considered the acclimation period and were discarded from the analysis. Mice had access to food and water ad libitum.

### Strength assay

Grip strength was assessed via a Cling assay on postoperative week 12. In brief, the time to loss of grip is collected for mice suspended on an overturned wire cage top. Four trials were conducted per mouse and averaged.

### Tissue harvest and RNA extraction

Mice were euthanized at 14 weeks post-operatively using cervical dislocation after a 16-hour fast and a 4-hour refeed. Tissues were flash-frozen in liquid nitrogen. Liver samples for RNA extraction were processed using the PureLink^TM^ RNA mini kit (Invitrogen; Carlsbad, CA) and submitted for transcriptomics.

### Construction of Stranded RNA Seq Libraries

The UW-Madison Biotechnology Gene Expression Center (Research Resource Identifier - RRID:SCR_017757) was utilized for RNA library preparation and the DNA Sequencing Facility (RRID:SCR_017759) for sequencing. Each submitted RNA sample was assayed on NanoDrop One Spectrophotometer and Agilent 4200Tapestaion to assess purity and integrity. RNA samples that met Illumina’s “TruSeq Stranded mRNA Reference Guide” (Document# 1000000040498 v00, October 2017) quality criteria were prepared for sequencing. Briefly, 1ug total RNA sample was enriched for mRNA by oligo (dT) selection, followed by fragmentation using divalent cations under elevated temperature. Double-stranded cDNA was synthesized using SuperScript II RT (Invitrogen, Carlsbad, CA) and random primers for first strand cDNA synthesis followed by second strand synthesis using DNA Polymerase I and RNAse H for removal of mRNA. Double-stranded cDNA was purified with SPRI beads (Beckman Coulter Genomics, San Jose, CA), followed by incubation with Klenow DNA Polymerase and Adenine to add an ‘A’ base (Adenine) to the 3’ end of the blunt phosphorylated DNA fragments. DNA fragments were ligated to Illumina unique dual index (UDI) adapters, which have a single ‘T’ base (Thymine) overhang at their 3’end. Following ligation, cDNA was purified 2x with SPRI beads, and PCR-amplified (11 cycles) with Phusion^TM^ DNA Polymerase followed by a final purification with SPRI beads.

Quality and quantity of the finished libraries were assayed on Agilent 4200Tapestation using D1000 Screentape and Agilent BioMek Synergy H1 Plate Reader with Picogreen reagent, respectively.

### Illumina Sequencing

Paired end 150bp sequencing was carried out on an Illumina NovaSeqX Plus sequencer with libraries multiplexed across six lanes on a 10B flowcell.

### Metabolomics

Hepatic tissue samples were ground at liquid nitrogen temperature with a CryoMill (Retsch). The resulting tissue powder (approximately 20 mg) was weighed and then mixed with −20°C extraction solvent containing 0.5% formic acid (40 µl/ mg tissue), vortexed and added 15% NH4HCO3(3.5µl/ mg tissue) to neutralize the acid. Following vortexing and centrifugation at 16,000g for 10 minutes at 4°C, the supernatant (70µl) was loaded to individual LC–MS vials. Metabolites were analyzed by quadrupole-orbitrap mass spectrometers (Q-Exactive Plus Hybrid Quadrupole-Orbitrap, Thermo Fisher) coupled to hydrophilic interaction chromatography (HILIC) via electrospray ionization. LC separation was on an Xbridge BEH amide column (2.1 mm x 150 mm, 2.5 µm particle size, 130 Å pore size; Waters) at 25°C using a gradient of solvent A (5% acetonitrile in water with 20 mM ammonium acetate and 20 mM ammonium hydroxide) and solvent B (100% acetonitrile). The flow rate was 150 µL/min. The LC gradient was: 0 min, 90% B; 2 min, 90% B; 3 min, 75% B; 7 min, 75% B; 8 min, 70% B; 9 min, 70% B; 10 min, 50% B; 12 min, 50% B; 13 min, 25% B; 14 min, 20% B; 15 min, 20% B; 16 min, 0% B; 20.5 min, 0% B; 21 min, 90% B; 25 min, 90% B. For bile acid analysis, 10 µL of blood were extracted with 300 µL ice-cold acetonitrile:methanol:water (40:40:20) solution.

Following vortexing and centrifugation, 70uL of supernatant was loaded to MS vials. Metabolites were analyzed by quadrupole-orbitrap mass spectrometers (Thermo Fisher) via electrospray ionization. LC separation was on an Atlantis T3 Column (150 x 2.1mm, 3 μm particle size; Waters) at 25°C using a gradient of solvent A (5% acetonitrile in water with 20 mM ammonium acetate and 20 mM ammonium hydroxide) and solvent B (100% acetonitrile). The flow rate was 150 µL/min. The LC gradient was: 0 min, 90% B; 2 min, 90% B; 3 min, 75% B; 7 min, 75% B; 8 min, 70% B; 9 min, 70% B; 10 min, 50% B; 12 min, 50% B; 13 min, 25% B; 14 min, 20% B; 15 min, 20% B; 16 min, 0% B; 20.5 min, 0% B; 21 min, 90% B; 25 min, 90% B.

Autosampler temperature was set at 4°C and the injection volume of the sample was 3 μL. MS data were acquired in negative and positive ion mode with a full-scan mode from m/z 70 to 830 and 140,000 resolution. Data were analyzed using the Compound Discoverer (Thermo Fisher Scientific) and the MAVEN software (build 682, http://maven.princeton.edu/index.php). Natural isotope correction for dual isotopes was performed with AccuCor2 R code (https://github.com/wangyujue23/AccuCor2) and IsoCorrectoR.^83^

## Data Integration

WGCNA analysis was conducted in R (Version 4.4.3) using the WGCNA package. First, we filtered by gene expression variance, keeping the top 50% variable genes to remove “noisy” genes; this reduced our gene number from 15523 to 7762 genes. We then checked that all genes had enough samples and there were no clear outliers before running the analysis. WGCNA analysis identifies significant modules of genes and their associations with phenotypes. Once modules of genes were identified, they were enriched for GO Pathways. We identified genes that have a high significance for protein intake and EE as well as high module membership in interesting modules.

For the differential expression heatmap, three data types (metabolomics, transcriptomics, and phenotypic outcomes) were used. To identify molecules of interest in each of the-omics datasets, significantly differentially expressed molecules between Sham vs SG in each protein group were used. These selected data, in addition to phenotypic data from SG animals were log2 transformed. To integrate the data, all three data types were concatenated. Correlations were performed using Spearman’s rank (1607 x 1607 = 2582449 correlations). Complete hierarchical clustering was used to reorder molecules based on 1 – Spearman correlation between all molecules. The number of clusters were determined by silhouette scores.^40^ Once modules were identified, metabolite enrichment in Metaboanalyst and gene enrichment in KEGG were used to generate pathways.

### Statistics

GraphPad Prism, version 10 (GraphPad Software, San Diego, CA, USA) was used for data analysis. Students’ t tests, general linear modeling, one-way ANOVA with Dunnet corrections, and area under-the-curve analyses were used where appropriate. Given that WD animals were not fed an isocaloric diet, Sham and SG animals in this diet group were compared separately (Welch’s T-test) from the protein groups (two-factor ANOVA with multiple comparisons). Omics data analysis was conducted in R (Version 4.3.3.).

## Supplemental Figures

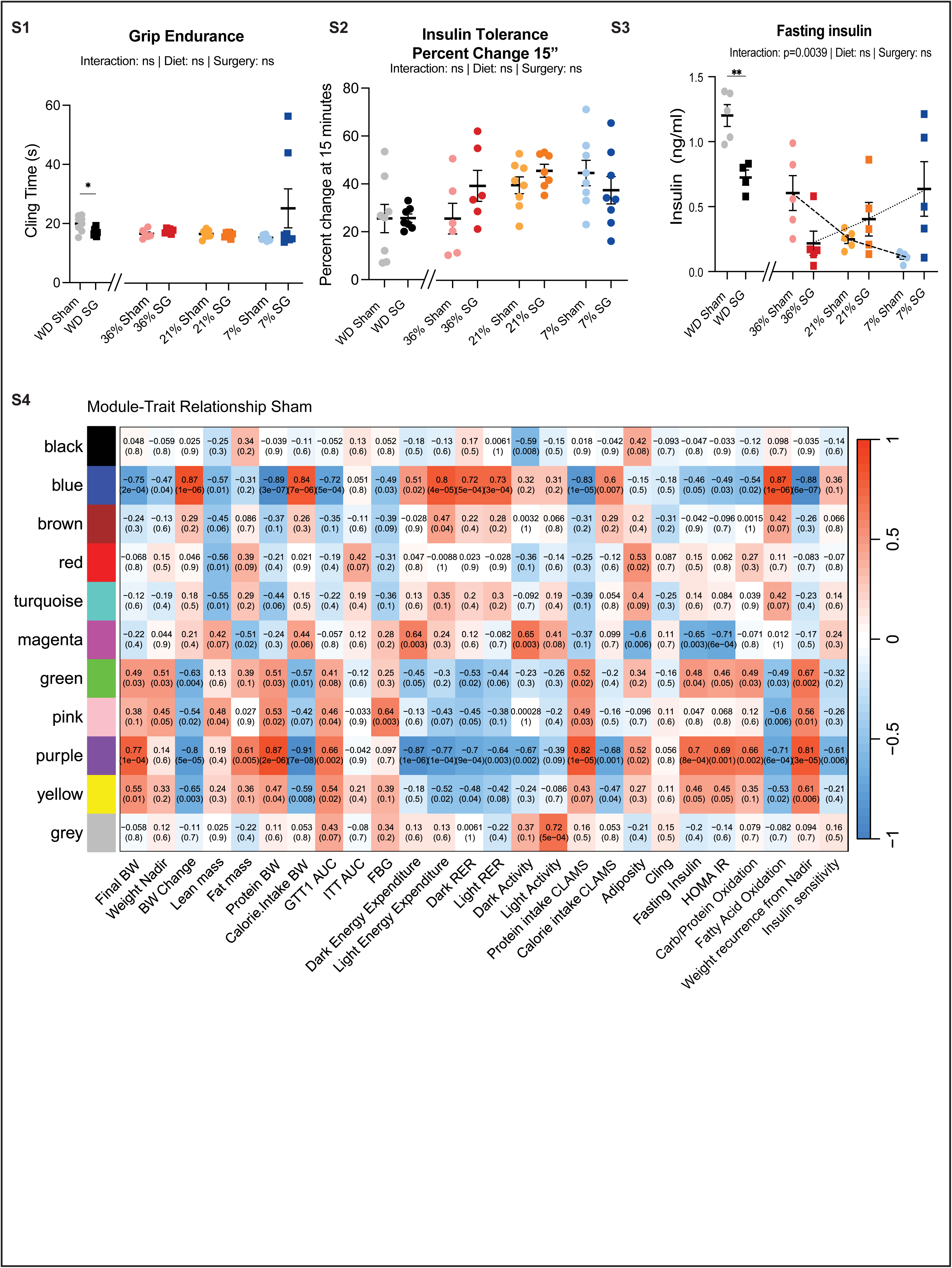
(**S1**) Inverted Cling assay (**S2**) Percent change at 15 minutes during insulin tolerance testing (**S3**) Fasting serum insulin (**S4**) WGCNA analysis of the Sham mice (**S1-S3**) n for each measurement is detailed in Table S9. WD surgical groups were compared using Welch’s T-test and surgical groups within the protein dietary groups were compared using two-way ANOVA between surgery and diet with post-hoc Tukey corrected test for pairwise comparisons. *p<0.05, **p<0.01; ***p<0.001; ****p<0.0001.

