## Supplemental Tables for "A low protein diet drives short-and long-term improvements in metabolic health in a mouse model of sleeve gastrectomy"

Table S1. Diet composition

| **Natural Source Diets** | **Western Diet** | **High protein** | **Control** | **Low Protein** |
|---|---|---|---|---|
| Teklad Diet Name | 42% Fat | 36% Protein Calories | 21% Protein Calories | 7% Protein Calories |
| Teklad Diet Number | TD.88137 | TD.220289 | TD.180161 | TD.10192 |
| Color | | | Red | Blue |
| **Formula** | **g/KG** | **g/kg** | **g/kg** | **g/kg** |
| Corn | 0 | 430 | 430 | 430 |
| Casein | 195 | 0 | 0 | 0 |
| Whey Protein Isolate | 210 | 215 | 170 | 35 |
| DL-Methionine | 3 | 2 | 2 | 0.4 |
| Corn Starch | 150 | 6 | 150.8 | 287.4 |
| Maltodextrin | 0 | 100 | 100 | 100 |
| Corn Oil | 0 | 32 | 32 | 32 |
| Olive Oil | 0 | 32 | 32 | 32 |
| Cellulose | 50 | 30 | 30 | 30 |
| Sucrose | 341.46 | 0 | 0 | 0 |
| Cholesterol | 50 | 0 | 0 | 0 |
| Mineral Mix, AIN-93G-MX (94046) | 35 | 35 | 35 | 35 |
| Calcium Phosphate, dibasic | 4 | 8 | 8 | 8 |
| Vitamin Mix, Teklad (40060) | 10 | 10 | 10 | 10 |
| **% kcal from** | | | | |
| Protein | 15.2 | 36 | 21.2 | 7.1 |
| Carbohydrate | 42.7 | 44 | 58.8 | 73 |
| Fat | 42 | 19.9 | 20 | 20 |
| Kcal/g | 4.5 | 3.6 | 3.6 | 3.6 |
| **Amino Acid Profile** | | **g/kg** | **g/kg** | **g/kg** |
| Lysine | | 33.7 | 18.6 | 4.5 |
| Methionine | | 9.7 | 6.5 | 1.9 |
| Cystine | | 12.3 | 7 | 2 |
| Arginine | | 10.8 | 6.8 | 3 |
| Phenylalanine | | 12.4 | 7.4 | 2.6 |
| Tyrosine | | 13.5 | 7.9 | 2.6 |
| Histidine | | 6.6 | 4 | 1.6 |
| Isoleucine | | 19.7 | 11.3 | 3.5 |
| Leucine | | 46.6 | 27.3 | 9.4 |
| Threonine | | 17.2 | 10.1 | 3.4 |
| Tryptophan | | 7.4 | 4.2 | 1.2 |
| Valine | | 18.3 | 10.7 | 3.7 |
| Aspartic Acid | | 39.2 | 22.4 | 6.7 |
| Glutamic Acid | | 52.3 | 28.6 | 6.4 |
| Alanine | | 19.5 | 11.8 | 4.7 |
| Glycine | | 6.9 | 4.5 | 2.2 |
| Proline | | 17.4 | 11.4 | 5.7 |
| Serine | | 15.3 | 9.2 | 3.6 |
| **Totals** | | 358.8 | 209.7 | 68.7 |

Table S2. Comparison of energy expenditure by lean mass across Sham groups by linear regression modeling.

| Diet Comparison | | Slope | | Y-Intercept | |
|---|---|---|---|---|---|
| **Diet 1** | **Diet 2** | **P-Value** | **Significance** | **P-Value** | **Significance** |
| **7% Sham** | **36% Sham** | **0.8858** | **No** | **0.0082** | **Yes** |
| 36% Sham | 21% Sham | 0.4047 | No | 0.2323 | No |
| 21% Sham | 7% Sham | 0.8251 | No | 0.0810 | No |
| **WD Sham** | **36% Sham** | **0.1406** | **No** | **0.0002** | **Yes** |
| **WD Sham** | **21% Sham** | **0.2574** | **No** | **<0.0001** | **Yes** |
| **WD Sham** | **7% Sham** | **0.4269** | **No** | **0.0009** | **Yes** |

Table S3. Comparison of energy expenditure by lean mass across SG groups by linear regression modeling.

| Diet Comparison | | Slope | | Y-Intercept | |
|---|---|---|---|---|---|
| **Diet 1** | **Diet 2** | **P-Value** | **Significance** | **P-Value** | **Significance** |
| **7% SG** | **36% SG** | **0.1818** | **No** | **0.0449** | **Yes** |
| 36% SG | 21% SG | 0.4540 | No | 0.2062 | No |
| 21% SG | 7% SG | 0.6227 | No | 0.2040 | No |
| WD SG | 36% SG | 0.2644 | No | 0.0676 | No |
| WD SG | 21% SG | 0.4486 | No | 0.0682 | No |
| WD SG | 7% SG | 0.5888 | No | 0.1122 | No |

Table S4. Top Ten Gene Changes Sham vs. SG – 21%

| Downregulated | | | Upregulated | | |
|---|---|---|---|---|---|
| **Gene** | **Fold Change** | **Adjusted p-value** | **Gene** | **Fold Change** | **Adjusted p-value** |
| Gm14584 | -1.93 | 0.024 | Gm20547 | 7.23 | 0.027 |
| Zfp771 | -1.88 | 0.013 | Orm2 | 2.82 | 0.038 |
| Gm56874 | -1.73 | 0.024 | Orm3 | 2.56 | 0.044 |
| Atp6v0c-ps2 | -1.73 | 0.018 | Tex14 | 2.54 | 0.013 |
| Rn7s2 | -1.67 | 0.026 | BB218582 | 2.53 | 0.016 |
| Rn7s1 | -1.67 | 0.026 | Gm37033 | 2.25 | 0.012 |
| Gm12715 | -1.65 | 0.017 | Cxcl2 | 2.23 | 0.041 |
| Lamb3 | -1.61 | 0.010 | Ttc39aos1 | 2.18 | 0.041 |
| AA465934 | -1.57 | 0.017 | Gm53048 | 2.16 | 0.014 |
| Csad | -1.56 | 0.001 | Gm34084 | 2.14 | 0.026 |
| Snx29 | -1.53 | 0.009 | Klrb1b | 2.11 | 0.029 |
| Gm57138 | -1.52 | 0.002 | Igkv15-103 | 2.09 | 0.038 |
| Oaz1-ps | -1.52 | 0.010 | Mup-ps16_1 | 2.08 | 0.021 |
| Gm11586 | -1.51 | 0.044 | Mefv | 2.06 | 0.040 |
| Marcksl1-ps4 | -1.44 | 0.040 | Sirpb1b | 1.93 | 0.015 |

Table S5. Top Gene Changes Sham vs. SG – 36%

| Down regulated | | | Upregulated | | |
|---|---|---|---|---|---|
| **Gene** | **Fold Change** | **Adjusted p-value** | **Gene** | **Fold Change** | **Adjusted p-value** |
| A1bg | -3.95 | 0.020 | Scara5 | 4.49 | 0.019 |
| Acot1 | -3.30 | 0.021 | Cyp21a1 | 3.83 | 0.045 |
| Ankrd40cl | -2.72 | 0.031 | Gm15348 | 3.82 | 0.049 |
| Dbp | -2.53 | 0.000 | Asns | 3.57 | 0.027 |
| Cidec | -2.52 | 0.046 | Fmn2 | 3.38 | 0.011 |
| Fam89a | -2.33 | 0.006 | Pnpla3 | 3.14 | 0.006 |

| | | | | | |
|---|---|---|---|---|---|
| Cyp4a10 | -2.27 | 0.001 | Hapln1 | 3.05 | 0.002 |
| Vnn1 | -2.26 | 0.000 | Meg3 | 3.05 | 0.001 |
| Cyp4a14 | -2.22 | 0.022 | Mirg | 3.03 | 0.045 |
| Gm22146 | -2.20 | 0.045 | Orm3 | 2.75 | 0.046 |
| 1810053B23Rik | -2.11 | 0.011 | Enho | 2.67 | 0.021 |
| Osbpl3 | -2.03 | 0.022 | Rian | 2.66 | 0.002 |
| Nr1d1 | -1.99 | 0.000 | Pex5l | 2.63 | 0.047 |
| 1810044K17Rik | -1.96 | 0.025 | Fam81a | 2.61 | 0.004 |
| Abcd2 | -1.95 | 0.020 | Chrm1 | 2.47 | 0.021 |

Table S6. Sham vs. SG – Shared significant gene changes 21 and 36%

| Gene | Fold Change 21% | Fold Change 36% |
|---|---|---|
| 2010320M18Rik | -0.93 | 0.005326478 |
| Rtf2 | -0.52 | 0.024460768 |
| **Hsdl2** | **-1.09** | **-0.257636087** |
| **Gan** | **1.22** | **0.221775008** |
| **Acadm** | **-0.86** | **-0.158444753** |
| Gpd1 | -1.30 | 0.032603272 |
| Acer2 | 0.70 | -0.104679557 |
| **Orm3** | **2.75** | **2.003777807** |
| **Gm33543** | **1.34** | **0.18749615** |
| **Csad** | **-1.85** | **-0.382868522** |
| **Gabarapl1** | **-1.01** | **-0.005075365** |
| **Etfdh** | **-0.88** | **-0.162552067** |
| **Nfkbiz** | **1.20** | **0.516171609** |
| **Isg15** | **-1.39** | **-0.045399101** |
| Commd9 | -0.60 | 0.095794969 |
| Sh3bp5l | -0.55 | 0.15612954 |
| **Irf9** | **-0.63** | **-0.121457511** |
| **Gm8355** | **-1.12** | **-0.015041164** |
| **Usp18** | **-1.38** | **-0.207240995** |
| Gm48878 | 1.418881373 | -0.289767689 |
| **Stat1** | **-0.812234333** | **-0.147331831** |
| **Crat** | **-1.690911457** | **-0.06434453** |
| Ctdsp1 | -0.439646446 | 0.060018587 |
| **Bsdc1** | **-0.558962543** | **-0.002087564** |
| Arsa | -0.709276593 | 0.051644399 |

Table S7. Top Gene Changes Sham vs. SG – 7%

| Gene | Fold Change | Adjusted p-value |
|---|---|---|
| Krt23 | -1.89 | 0.007 |

Table S8. Top 10 up and down regulated gene changes – 7 vs 36%SG

| Gene | Fold Change | Adjusted p-value |
|---|---|---|
| FGF21 | -6.5193348 | 0.01373738 |
| Cps1 | -5.8286214 | 0.00129799 |

| | | |
|---|---|---|
| Gm15998 | -4.7924899 | 0.00483949 |
| Gm51903 | -4.3599419 | 0.00047579 |
| Pmm2 | -4.2156478 | 0.00017441 |
| Lrrc39 | -4.204143 | 0.00013963 |
| Mtmr10 | -3.9966525 | 0.02727162 |
| Fut1 | -3.9878198 | 0.00028613 |
| Asb4 | -3.9786819 | 0.00492898 |
| Cfb | -3.9757187 | 0.00402499 |
| Zfp664 | 7.28770441 | 0.0221906 |
| Cyp21a1 | 5.30495178 | 0.00085582 |
| Lrtm2 | 5.07550869 | 0.00068207 |
| Chrna4 | 4.04649313 | 0.00237001 |
| Ctsd | 3.71810178 | 0.01090143 |
| Gm43802 | 3.6812388 | 0.02029596 |
| Gm15348 | 3.57011313 | 0.00473012 |
| Gm10762 | 3.45966108 | 0.00952928 |
| Tiparp | 3.42273575 | 0.00061377 |
| Orm2 | 3.42271024 | 0.01625033 |

Table S9. Number of animals represented in each figure.

| **Figures** | **Surgery Group** | **WD** | **36% PR** | **21% PR** | **7% PR** |
|---|---|---|---|---|---|
| 1B | Sham | 8 | 6-8 | 8 | 8 |
| | SG | 8 | 5-8 | 7-8 | 7-8 |
| 1C | Sham | 8 | 6-8 | 8 | 8 |
| | SG | 8 | 5-8 | 7-8 | 7-8 |
| 1D | Sham | -- | -- | -- | -- |
| | SG | 8 | 6 | 7 | 7 |
| 1E | Sham | 8 | 6-8 | 8 | 8 |
| | SG | 8 | 5-8 | 7-8 | 7-8 |
| 1F | Sham | 8 | 6 | 8 | 8 |
| | SG | 8 | 6 | 7 | 7 |
| 1G | Sham | 8 | 6 | 8 | 8 |
| | SG | 8 | 6 | 7 | 7 |
| 1H | Sham | 8 | 6 | 8 | 8 |
| | SG | 8 | 6 | 7 | 7 |
| 1I | Sham | -- | -- | -- | -- |
| | SG | 8 | 5 | 7 | 7 |
| 2A | Sham | 4* | 6 | 8 | 8 |
| | SG | 8 | 6 | 6* | 7 |
| 2B | Sham | 4* | 6 | 8 | 8 |
| | SG | 8 | 6 | 7 | 7 |
| 2C | Sham | 4* | 6 | 8 | 8 |
| | SG | 8 | 6 | 7 | 7 |
| 2D | Sham | 4* | 6 | 8 | 8 |
| | SG | 8 | 6 | 7 | 7 |
| 2E | Sham | 4* | 6 | 8 | 8 |
| | SG | 8 | 6 | 7 | 7 |

| | | | | | |
|---|---|---|---|---|---|
| 2F | Sham | 4* | 6 | 8 | 8 |
| | SG | 8 | 6 | 7 | 7 |
| 2G | Sham | 6 | -- | -- | 8 |
| | SG | -- | -- | -- | -- |
| 2H | Sham | -- | -- | -- | -- |
| | SG | 6 | -- | -- | 7 |
| 2I | Sham | 4* | 6 | 8 | 8 |
| | SG | 8 | 6 | 7 | 7 |
| 2J | Sham | 4* | 6 | 8 | 8 |
| | SG | 8 | 6 | 7 | 7 |
| 3A | Sham | 8 | 6 | 8 | 8 |
| | SG | 8 | 8 | 7 | 8 |
| 3B | Sham | 8 | 6 | 8 | 8 |
| | SG | 8 | 8 | 7 | 8 |
| 3C | Sham | 8 | 6 | 8 | 8 |
| | SG | 8 | 8 | 7 | 8 |
| 3D | Sham | 8 | 6 | 8 | 8 |
| | SG | 7* | 6 | 7 | 8 |
| 3E | Sham | 8 | 6 | 8 | 8 |
| | SG | 7* | 6 | 7 | 8 |
| 3F | Sham | 5 | 5 | 5 | 5 |
| | SG | 4* | 5 | 5 | 5 |

*Outliers excluded using Grubb's test
